# Lymphatic dissemination is a major route of systemic invasion by diverse extracellular bacteria in soft tissue infection

**DOI:** 10.1101/2025.04.21.649860

**Authors:** Matthew K. Siggins, Ho Kwong Li, Kristin K. Huse, Max Pearson, Peter J.M. Openshaw, David G. Jackson, Shiranee Sriskandan

## Abstract

Extracellular bacterial pathogens cause substantial global mortality, yet how they disseminate from peripheral tissues into the bloodstream remains poorly understood. Systemic infection is typically attributed to direct vascular invasion, with draining lymph nodes considered terminal sites of containment. Here, using a murine soft-tissue infection model, we show that onward lymphatic dissemination from a distant tissue site through sequential draining lymph nodes is a major route of systemic invasion across diverse extracellular bacterial pathogens. Clinical isolates of *Escherichia coli, Klebsiella pneumoniae, Pseudomonas aeruginosa, Streptococcus pyogenes*, and *Streptococcus agalactiae* spread from local draining lymph nodes to downstream lymph nodes and systemic organs. Non-draining lymph nodes, exposed to the same systemic circulation as downstream draining lymph nodes, contained markedly fewer bacteria even during bacteraemia, indicating lymphatic rather than haematogenous spread. *Staphylococcus aureus* was a notable exception, remaining largely restricted to local draining nodes with limited systemic spread. Deletion of the *S. agalactiae* capsular polysaccharide significantly reduced lymphatic and systemic spread without significantly reducing bacterial burden at the infection site, showing that capsule can promote lymphatic dissemination independently of local bacterial burden. In contrast, digestion of the secreted exopolysaccharide of *P. aeruginosa* did not alter dissemination, distinguishing the impact of surface-attached capsule from secreted exopolysaccharide. These findings establish lymphatic dissemination through sequential lymph nodes as a major invasion route across key extracellular bacterial pathogens, with important implications for systemic disease and immunity.

## Introduction

Extracellular bacterial pathogens account for an estimated 6 million deaths annually and dominate antimicrobial resistance priority lists [1, 2], but incomplete understanding of systemic invasion constrains efforts to prevent, predict, and treat serious disease. More than a quarter of bacterial bloodstream infections lack a defined primary focus [3].

Lymphatic vessels permeate tissues throughout the body, returning interstitial fluid to the blood through lymph nodes central to adaptive immunity [4]. Lymphatic dissemination is recognised in cancer [5] and a small number of specialised intracellular lymph-tropic infections, including *Yersinia pestis*, which can transit through lymphatics within phagocytic cells [6]. Despite providing a direct anatomical route from peripheral tissues to the circulation, the lymphatic system remains poorly defined in extracellular bacterial infection [7, 8].

Lymph nodes are commonly assumed to contain and eliminate pathogens, preventing onward spread to sequential draining lymph nodes and the bloodstream [9, 10]. However, this perspective has largely been shaped by studies of viruses [9, 11, 12], while evidence from diverse extracellular bacterial pathogens remains lacking.

## Results

To test whether lymphatic dissemination is a common route of invasion, we analysed clinical isolates from six extracellular bacterial species, encompassing the leading causes of uncomplicated and complicated soft-tissue infection and associated bacteraemia [2, 13, 14], in a standardised murine hindlimb infection model. At 6 h after intramuscular infection, we quantified bacterial burdens at the hindlimb infection site, and in blood, spleen, liver, and draining and non-draining lymph nodes.

All isolates of *E. coli, K. pneumoniae, P. aeruginosa, S. agalactiae*, and *S. pyogenes* were consistently recovered from the local draining inguinal and iliac lymph nodes, the downstream draining axillary lymph node, blood, and systemic organs (Fig 1A–E). By contrast, non-draining brachial lymph nodes, exposed to the same systemic circulation as the downstream draining axillary node, contained markedly fewer bacteria even during high-grade bacteraemia, indicating lymphatic rather than haematogenous seeding. Although dissemination varied substantially between species and strains, bacterial burdens were significantly higher in draining than non-draining lymph nodes for all bacterial species. *S. aureus* was a notable exception: USA300 and contemporary clinical MRSA and MSSA isolates were largely confined to initial draining lymph nodes, with minimal onward spread to downstream lymph nodes or systemic sites (Fig 1F).

**Figure 1.**
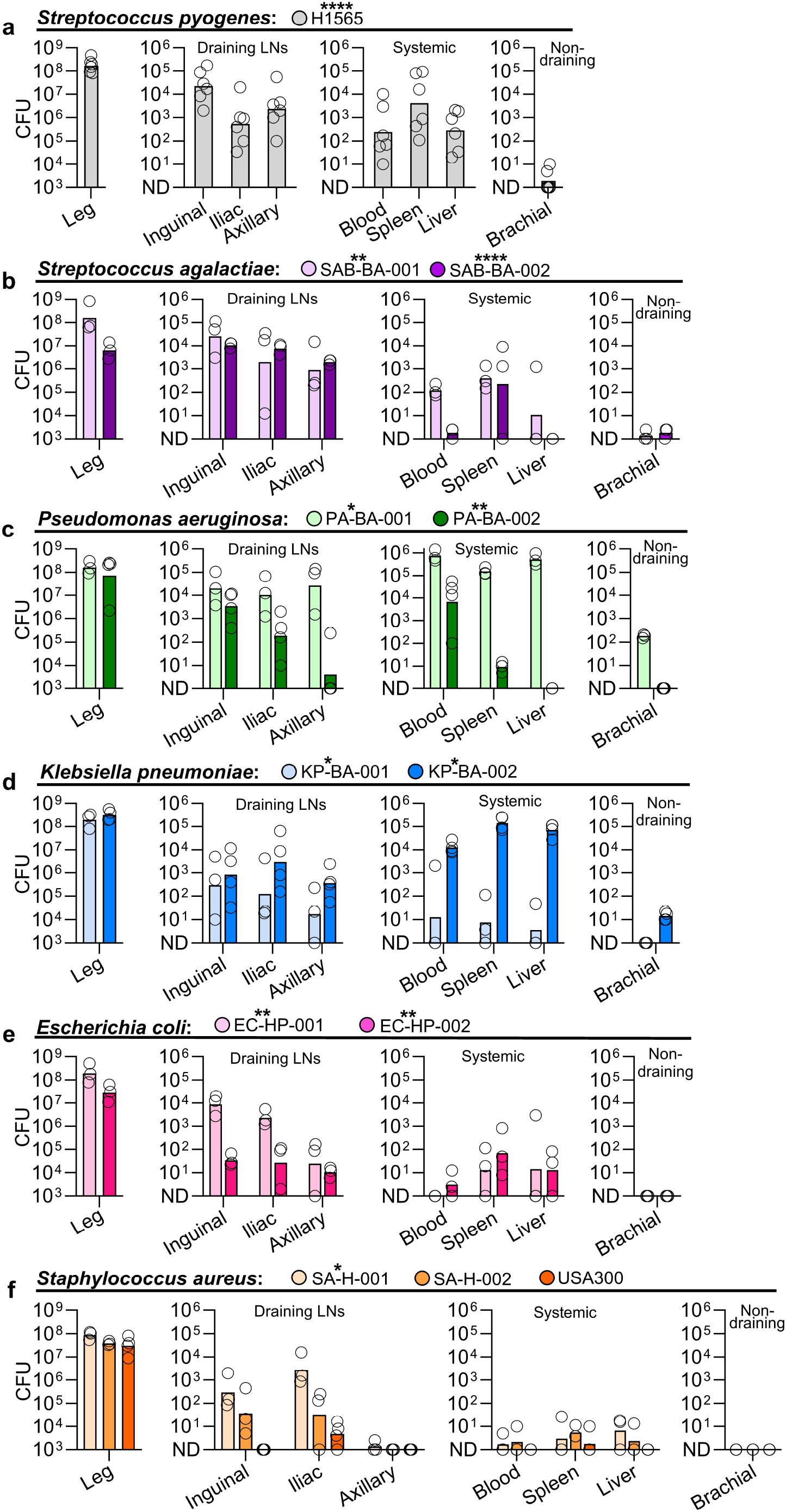
Diverse extracellular bacteria disseminate through sequential draining lymph nodes and to systemic organs. Recovery of *S. pyogenes* (a), *S. agalactiae* (b), *P. aeruginosa* (c), *K. pneumoniae* (d), *E. coli* (e), and *S. aureus* (f) from the infection site, draining and non-draining lymph nodes, blood, and systemic organs of FVB/n mice 6 h after intramuscular infection with 10^8^ CFU. Symbols represent individual mice (n = 4–6); bars indicate geometric means. CFU per ml blood, per gram of liver, per whole spleen, lymph node, and infected hindlimb tissue. Two-tailed Welch’s t-tests performed on log_10_-transformed data with Benjamini–Hochberg false discovery rate correction, comparing draining and non-draining nodes for each isolate. ****p ≤ 0.0001; **p ≤ 0.01; *p ≤ 0.05; ns, p > 0.05.

As capsular polysaccharide can alter bacterial surface properties and interactions with host tissues, we tested whether capsule promotes lymphatic dissemination. We compared encapsulated *S. agalactiae* A909 with its otherwise isogenic capsule-deficient Δ*cpsD* mutant. At 3 h after intramuscular infection, encapsulated *S. agalactiae* had disseminated to and through local and downstream draining lymph nodes to blood, spleen, and liver, while non-draining lymph nodes contained few bacteria. Loss of capsule significantly reduced bacterial burden in local and downstream draining lymph nodes as well as systemic organs (Fig 2A). Bacterial burdens at the infection site did not differ between strains, indicating that reduced lymphatic dissemination was not explained by reduced bacterial abundance at the initial infection site.

**Figure 2.**
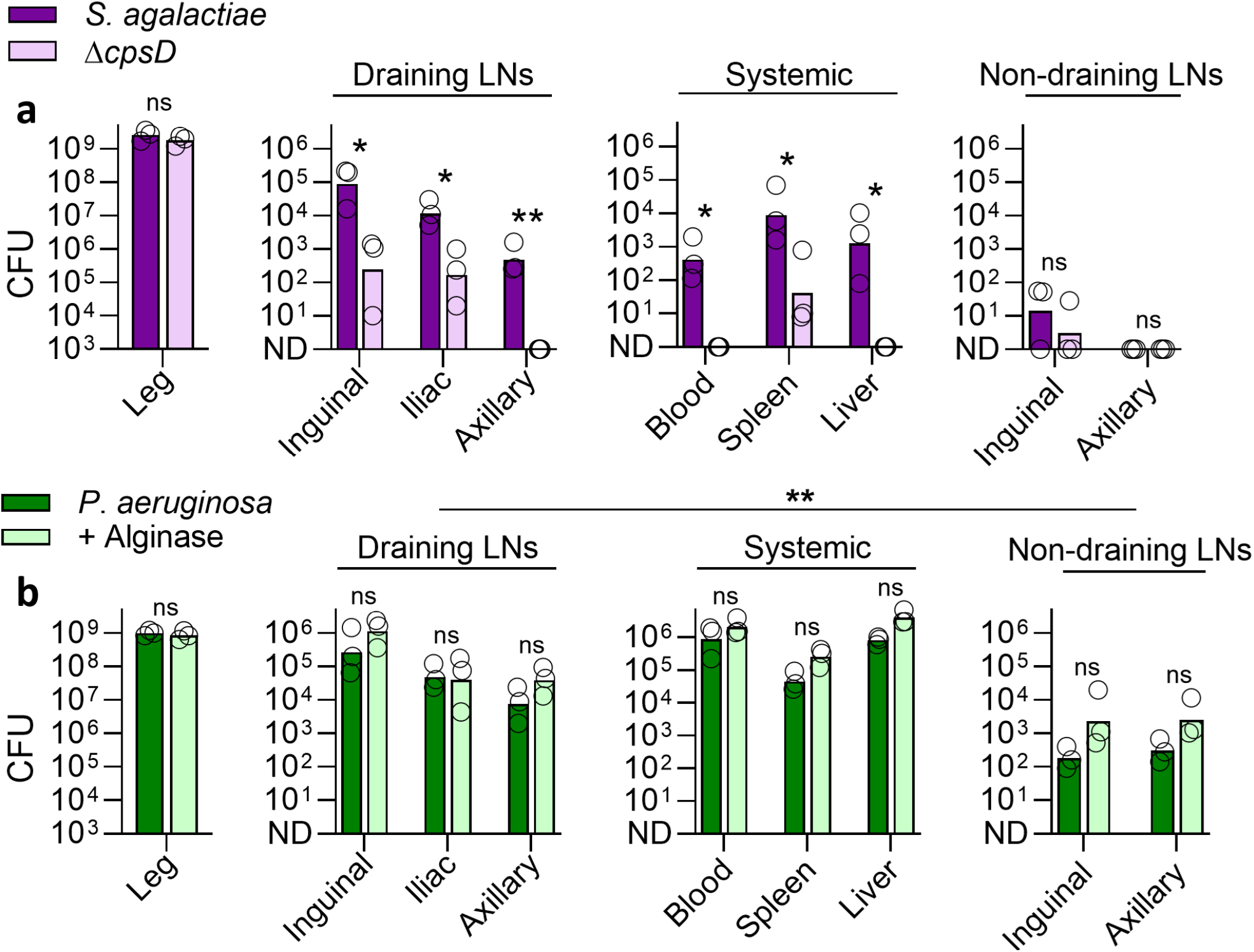
S. agalactiae capsular polysaccharide, but not *P. aeruginosa-* secreted alginate exopolysaccharide, promotes lymphatic dissemination. (a) Recovery of *S. agalactiae* A909 and its isogenic Δ*cpsD* mutant, and (b) of *P. aeruginosa* PA-BA-001 with or without alginate lyase pre-treatment, 3 h after intramuscular infection with 10^8^ CFU. Symbols represent individual mice (n = 4); bars indicate geometric means. Bacterial CFU expressed as reported in Figure 1. Two-tailed Welch’s t-tests performed on log_10_-transformed data with Benjamini–Hochberg false discovery rate correction. **p ≤ 0.01; *p ≤ 0.05; ns, p > 0.05.

*P. aeruginosa* showed substantial lymphatic dissemination during infection despite lacking classical capsular polysaccharide like *S. agalactiae*. Instead, it produces a secreted alginate exopolysaccharide. We therefore tested whether exopolysaccharide could similarly promote dissemination by pre-treating *P. aeruginosa* with alginate lyase or control before infection. At 3 h after intramuscular infection, digestion of the alginate exopolysaccharide did not alter dissemination. Treated and untreated groups showed similarly high bacterial burdens in draining lymph nodes and systemic organs and significantly fewer bacteria in non-draining lymph nodes (Fig 2B).

## Discussion

Here, we show that lymphatic dissemination from distant infected tissue through sequential draining lymph nodes is common across diverse extracellular bacterial pathogens spanning leading causes of uncomplicated and complicated skin and soft tissue infections and associated bacteraemia. These findings challenge the prevailing view that local draining lymph nodes function as terminal traps for extracellular bacteria. That perspective has been shaped by prominent viral studies [9-11], while a limited bacterial literature has centred on *S. aureus* USA300 and containment by the initial draining lymph node [15-17]. Clinically, bacterial lymphatic dissemination also provides an anatomical explanation for cryptic bacteraemia and severe systemic disease arising from seemingly limited peripheral infection [3].

Onward dissemination through sequential draining lymph nodes occurred across extracellular bacterial pathogens with distinct cell morphologies, surface structures, and virulence strategies. In these infections, non-draining lymph nodes had significantly fewer bacteria than draining lymph nodes even during bacteraemia. Non-draining brachial lymph nodes are exposed to the same systemic circulation as downstream axillary draining lymph nodes, indicating that lymphatic dissemination predominated over haematogenous seeding. By contrast, *S. aureus* USA300 and contemporary clinical MRSA and MSSA isolates were largely confined to the initial draining nodes. *S. aureus* therefore appears to be an important exception rather than a representative model of extracellular bacterial dissemination.

As identical 10^8^ CFU inocula resulted in markedly varied lymphatic dissemination between pathogen species and strains, lymphatic dissemination cannot be an artefact of high bacterial burdens. Indeed, we have observed sequential lymphatic dissemination from a distant tissue site with as few as 10^4^ CFU of *S. pyogenes* [18]. Irrespective, high inocula of approximately 10^8^ CFU are well-established in murine soft-tissue infection models across all the bacterial species investigated here [19-23]. Although no typical founding inoculum concentration has been established in human infection, human intradermal challenge demonstrated that high numbers of *S. aureus*, between 10^6^ and 10^7^ CFU, were required to generate a pus-forming infection [24]. High bacterial burdens are also well documented in human soft-tissue infections, averaging above 10^8^ CFU/ml in infected pus and peritoneal fluid [25], and exceeding 10^8^ CFU/g in a fifth of wound biopsies [26]. Clinical fluids obtained from invasive *S. pyogenes* infection contained a median of 10^9^ genome copies/ml, reaching as high as 5×10^11^ [27].

Lymphatic dissemination was not restricted to recognised lymph-tropic pathogens. We previously showed that extracellular lymphatic metastasis drives *S. pyogenes* systemic infection [18], but its clinical lymphatic association and hyaluronan capsule, which binds the lymphatic endothelial receptor LYVE-1, suggested this was a pathogen-specific adaptation [28]. The similar dissemination we have observed across unrelated bacteria shows that lymphatic spread is not a specialised feature of *S. pyogenes*, though the mechanisms driving it may vary.

Species and strain variation suggest that virulence factors influence lymphatic dissemination. Deletion of the *S. agalactiae* capsule markedly reduced lymphatic and systemic spread without significantly altering bacterial burden at the initial infection site, indicating a role beyond local survival. This is consistent with our finding that depletion of lymph node macrophages and neutrophils did not reduce *S. pyogenes* lymphatic dissemination [18]. Together, our data show structurally distinct surface-associated polysaccharide capsules can promote lymphatic dissemination. The polyanionic and hydrophilic properties of bacterial capsules may facilitate interstitial and lymphatic movement by masking adhesins and increasing electrostatic repulsion in the extracellular matrix [29-34]. However, for *S. aureus*, neither the acapsular USA300 nor the capsule-expressing MSSA (CC15) and MRSA (CC22) isolates reached the distant draining axillary lymph node, indicating that capsule presence alone does not explain onward lymphatic dissemination. Similarly, digestion of the secreted alginate exopolysaccharide of *P. aeruginosa* did not reduce lymphatic dissemination.

By delivering viable bacteria into collecting lymphatic vessels and draining lymph nodes that orchestrate immune responses, lymphatic dissemination may disrupt lymphatic function and immune organisation. Bacterial infection can disrupt macrophage and dendritic cell organisation, germinal centre formation, and T-cell responses [35-38] and *S. aureus* can impair lymphatic contractility and flow in pre-nodal vessels [39]. The breadth of lymphatic dissemination observed here suggests that bacterial–lymphatic interactions are more widespread than recognised and are an underexplored component of bacterial disease and immunity.

Our data establish lymphatic dissemination as a major invasion route across diverse extracellular bacterial pathogens. A full understanding of systemic bacterial infection and host responses must therefore incorporate lymphatic spread alongside haematogenous dissemination and bacterial replication within organs.

## Materials and Methods

### Bacterial strains and growth conditions

*Streptococcus pyogenes* and *Streptococcus agalactiae* were cultured on Columbia horse blood agar (CBA; EO Labs) or in Todd Hewitt broth supplemented with yeast extract (both Oxoid) at 37°C in 5% CO2. *Escherichia coli, Klebsiella pneumoniae, Pseudomonas aeruginosa* and *Staphylococcus aureus* were grown on CBA at 37°C in air overnight. All strains and their origins are described below (Bacterial strains used in this study). The Bioresource in Adult Infectious Diseases (BioAID)[40] and Microbial Products [41] are REC-approved bioresources of patient isolates. BioAID is a multicentre bioresource that collects biological samples from patients with suspected infection at presentation to the Emergency Department.

### Mice

In vivo experiments were performed in accordance with the Animals (Scientific Procedures) Act 1986, under appropriate UK Home Office licences and according to established institutional guidelines, using 6–8-week-old female FVB/n mice (Charles River UK).

### Infections

Mice were challenged intramuscularly with 10^8^ CFU of *S. pyogenes, S. agalactiae, E. coli, K. pneumoniae, P. aeruginosa*, or *S. aureus* in 50 µl PBS, and quantitative endpoints were compared at 3 h or 6 h post-infection as indicated. For alginate exopolysaccharide digestion, 200 U of alginate lyase was added to a *P. aeruginosa* inoculum and incubated for 30 minutes at 37°C prior to infection.

### Enumeration of bacteria from organs and blood

At the specified endpoints, mice were anaesthetised with isoflurane (4% induction, 2% maintenance), and blood was collected by cardiac puncture into tubes containing heparin before euthanasia. Thigh muscle, liver, spleen, and the ipsilateral axillary, brachial, iliac, and inguinal lymph nodes and contralateral inguinal and axillary nodes were dissected for analysis. Dissected lymph nodes were first disrupted with a motorised pellet pestle, and all organs and tissues were then homogenised with scissors in PBS. Bacterial CFU were determined by plating homogenised tissue and blood samples onto the specified agar, with or without dilution in PBS as appropriate. CFU are displayed per ml of blood, per g of liver, per leg, or per organ for all figures.

### Statistics

All statistical analyses were performed with GraphPad Prism 11. Comparison of two datasets was carried out using two-tailed Welch’s t-tests on log_10_-transformed data, with p-values adjusted for multiple comparisons using the Benjamini–Hochberg false discovery rate method. Comparisons of groups of draining lymph nodes and groups of non-draining lymph nodes normalised CFU counts by the number of lymph nodes per group.

### Study approval

Use of clinical isolates was approved by the South Central – Oxford C Research Ethics Committee (14/SC/0008 and 19/SC/0116; BioAID) and the West London Research Ethics Committee (06/Q0406/20; NIHR Health Protection Research Unit). All animal experiments were performed in accordance with the Animals (Scientific Procedures) Act 1986 under UK Home Office project licence.

### Bacterial strains

Twelve clinical isolates spanning six species, together with reference and mutant strains, were used; known typing, clinical source, and bioresource identifiers are summarised below: *Escherichia coli*: two bacteraemia isolates from the Microbial Products bioresource: EC-HP-001 (sequence type ST131) and EC-HP-002 (ST88).

*Klebsiella pneumoniae*: two hypermucoviscous (string test–positive) isolates from BioAID: KP-BA-001 (H1733), of undefined type, and KP-BA-002 (H1521), of undefined type, originating from a skin infection.

*Pseudomonas aeruginosa*: two isolates from BioAID, both of undefined type: PA-BA-001 (H1730), from a bacteraemia, and PA-BA-002 (H1522), from a breast abscess.

*Streptococcus agalactiae*: two neonatal bacteraemia isolates from BioAID, SAB-BA-001 (H1146; serotype Ia) and SAB-BA-002 (H1148; serotype V), and the serotype Ia reference strain A909 (a 1934 neonatal bacteraemia isolate) and its otherwise isogenic acapsular mutant, A909 Δ*cpsD*, constructed with pJRS233 (Areschoug and Lindahl, unpublished).

*Staphylococcus aureus*: three isolates: SA-H-001 (clonal complex CC15), a methicillin-sensitive (MSSA) bacteraemia isolate (“HHS-8”); SA-H-002 (CC22), a methicillin-resistant (MRSA) bacteraemia isolate (“HHS-9”); and the widely used community-acquired MRSA reference strain USA300 (CC8), which is naturally acapsular.

*Streptococcus pyogenes*: H1565, an encapsulated *covR* mutant derivative of the *emm1* maternal sepsis isolate H584, recovered after in vivo passage.

Clinical isolates were obtained from the BioAID and Microbial Products patient bioresources.

## Acknowledgements

We thank Alex McCarthy and Gunnar Lindahl for *S. agalactiae* A909 and its Δ*cpsD* mutant. The Imperial College NIHR Biomedical Research Centre supported the Leonard and Dora Colebrook Laboratory Infection Bioresource. Work was funded by MRC grant (MR/L008610/1) to S.S. and D.G.J. M.K.S. was supported in part by an NHLI Fellowship.

## Author contributions

Conceptualization: M.K.S., S.S.; Data curation: M.K.S., H.K.L., S.S.; Formal analysis: M.K.S.; Funding acquisition: M.K.S., D.G.J., S.S.; Investigation: M.K.S., M.P., K.K.H.; Methodology: M.K.S.; Resources: H.K.L., S.S.; Supervision: M.K.S., P.J.M.O., S.S.; Visualization: M.K.S.; Writing – original draft: M.K.S. **comparing draining and non-draining nodes for each isolate**

